# Hepatic ketogenesis supports liver lipid homeostasis during acute exercise but is not required for exercise training to mitigate liver steatosis in mice

**DOI:** 10.64898/2026.01.24.701392

**Authors:** Cha Mee Vang, Andres F. Ortega, Ruth E. Pfeiffer, Jared L. Hartmann, Griffin S. Hampton, Hu Wang, Eric D. Queathem, Peter A. Crawford, Xianlin Han, Curtis C. Hughey

**Author notes:** Correspondence: Curtis C. Hughey from the Department of Medicine, Division of Molecular Medicine, University of Minnesota, Dwan 316, 425 East River Parkway, Minneapolis, MN, 55455.

## Abstract

The acceleration of hepatic lipid disposal during acute exercise has been proposed as a contributor to the anti-steatotic effects of exercise training. Ketogenesis, which produces acetoacetate (AcAc) and β-hydroxybutyrate (βOHB) from fatty acids, is among the lipid disposal pathways stimulated by exercise. This study tested the hypothesis that hepatic ketogenesis is necessary for exercise training to lower liver lipids. Liver-specific 3-hydroxymethylglutaryl-CoA synthase 2 knockout (HMGCS2 KO) mice and wild type (WT) littermates underwent sedentary, acute exercise, and exercise training protocols. Liver ketone bodies and lipids were determined via mass spectrometry platforms. Stable isotope infusions in conscious, unrestrained mice defined mitochondrial oxidative fluxes at rest and during exercise. Loss of hepatic HMGCS2 decreased liver AcAc and βOHB concentrations and impaired their increase during exercise.

Liver triacylglycerides (TAGs) were comparable between genotypes at rest (i.e., ad libitum fed and short fasted conditions). In contrast, liver TAGs were elevated in HMGCS2 KO mice following acute, non-exhaustive exercise. Liver TCA cycle flux was higher in KO mice at rest. During exercise, TCA cycle flux increased in both WT and KO mice but was not different between genotypes with greater exercise duration. This suggests that enhanced disposal of lipids via the TCA cycle may prevent liver lipid accumulation in HMGCS2 KO mice under sedentary conditions, but not during exercise. Unexpectedly, exercise training decreased liver TAGs similarly in both HMGCS2 KO and WT mice. In conclusion, hepatic ketogenesis supports liver lipid homeostasis during acute exercise, but is not required for exercise training to lower liver lipids.

**NEW & NOTEWORTHY:** Exercise training has been proposed to mitigate liver steatosis partly through enhanced hepatic lipid disposal. During acute exercise, the disposal of fatty acids to ketone bodies is stimulated. This study tested the hypothesis that hepatic ketogenesis was required for exercise training to reduce liver fat in mice. The results show that hepatic ketogenesis is needed to prevent lipid accumulation during acute exercise, but is not necessary for exercise training to lower liver lipids.

## INTRODUCTION

Regular exercise is a first-line intervention for metabolic dysfunction-associated steatotic liver disease (MASLD) (1). The effectiveness of exercise to mitigate liver steatosis has been linked to its ability to enhance hepatic lipid disposal both acutely and chronically (2). During acute exercise, hepatic lipid disposal is functionally coupled to glucose production. More specifically, liver glucose production is increased to match the heightened glucose demands of muscular work (3, 4). The enhanced liver glucose production is accomplished partly by a rise in gluconeogenesis, an energetically costly process that contributes to the greater hydrolysis of ATP to ADP and AMP during exercise (3–9). To blunt this energy discharge, fatty acid catabolism via terminal oxidation (i.e., TCA cycle flux and oxidative phosphorylation) is stimulated (2, 7, 8).

Thus, exercise accelerates fat oxidation so that the potential energy in fatty acids is transformed to ATP to support gluconeogenesis and, subsequently, working muscle (10). We previously tested if the exercise-mediated elevation in fat disposal via terminal oxidation was required for exercise training to lower liver lipids. Loss of hepatic phosphoenolpyruvate carboxykinase 1 (PCK1) prevented the rise in gluconeogenic and TCA cycle flux during acute exercise in mice (8). However, the lowering of liver triacylglycerides (TAGs) by training was not impaired (8).

In addition to terminal oxidation, acute exercise engages other hepatic lipid disposal pathways responsible for transforming the potential energy in fatty acids to alternative forms of energy currency that are then released from the liver to support muscular work. This includes promoting the production of acetoacetate (AcAc) and β-hydroxybutyrate (βOHB) from fatty acids via ketogenesis (11). Ketogenesis is a mitochondrial pathway initiated by 3-hydroxymethylglutaryl-CoA synthase 2 (HMGCS2), which synthesizes HMG-CoA from β-oxidation-derived acetoacetyl-CoA and acetyl-CoA (12). Next, HMG-CoA lyase (HMGCL) cleaves HMG-CoA to produce AcAc (12). AcAc can be released into the circulation, however, most is reduced by βOHB dehydrogenase 1 (BDH1) to βOHB prior to hepatic output (11, 12). Exercise provokes ketogenesis by increasing fatty acid delivery to the liver, elevating hepatic extraction of fatty acids from the circulation, and enhancing the intrahepatic conversion of fatty acids to ketone bodies (4, 13). While these ketogenic responses to exercise have received significant investigation, it remains unclear whether the stimulation of ketogenesis during acute exercise contributes to the efficacy of exercise training to combat liver steatosis.

The experiments presented herein tested the hypothesis that hepatic ketogenesis is necessary for exercise training to lower liver lipids. Liver-specific HMGCS2 knockout (KO) mice and wild type (WT) littermates were fed a Gubra Amylin NASH (GAN) diet to promote fatty liver. WT and liver-specific HMGCS2 KO mice were untrained or exercise-trained via treadmill running. Targeted metabolomics was completed to quantify liver ketone bodies, shotgun lipidomics was performed to determine liver lipids, indirect calorimetry assessed whole-body energy metabolism, and stable isotope infusions in conscious unrestrained mice defined glucose and mitochondrial oxidative fluxes at rest and during exercise. The results of this study show that hepatic ketogenesis promotes liver lipid homeostasis during acute exercise, but is not necessary for exercise training to mitigate diet-induced fatty liver.

## MATERIALS AND METHODS

### Mouse models and husbandry

The University of Minnesota Institutional Animal Care and Use Committee approved all experimental procedures. Mice on a C57BL/6NJ background with a hepatocyte-specific KO of HMGCS2 were generated by breeding mice expressing Cre recombinase under control of the albumin promoter (B6.Cg-Tg(Alb-cre)21Mgn/J) with HMGCS2 floxed mice as previously described (14). Floxed HMGCS2 littermates of the liver-specific HMGCS2 KO mice that did not express Cre recombinase were used as WT controls. All mice were fed Teklad Global 18% Protein Rodent Diet (18% kcal from fat, 58% kcal from carbohydrate, and 24% kcal from protein; 2918, Inotiv, Indianapolis, IN) upon weaning at three weeks of age. At six weeks of age, mice were provided a Gubra Amylin NASH (GAN) diet (40% kcal from fat [mostly palm oil], 20% kcal from fructose, and 2% cholesterol; D09100310; Research Diets Inc., New Brunswick, NJ). WT and liver HMGCS2 KO mice remained on the GAN diet for 6, 12, or 32 weeks until they were euthanized at 12, 18, or 38 weeks of age. One cohort of WT and liver HMGCS2 KO mice received a ketogenic diet (90.5% kcal from fat and 9.5% kcal from protein; TD.160153, Inotiv, Indianapolis, IN) for 72 hours prior to being euthanized at 12 weeks of age. Additional studies used hepatocyte-specific BDH1 KO mice on a C57BL/6NJ background that were generated by crossing B6.Cg-Tg(Alb-cre)21Mgn/J mice with BDH1 floxed mice as previously described (15). Cre-negative, floxed BDH1 littermates were used as WT controls. These mice were fed the Teklad Global 18% Protein Rodent Diet (2918, Inotiv, Indianapolis, IN) from weaning until euthanized at 12 weeks of age. Deletion of liver HMGCS2 and BDH1 was confirmed by both PCR genotyping and immunoblotting. One cohort of 16-week-old C57BL/6J mice (Strain# 000664, The Jackson Laboratory, Bar Harbor, ME) fed the Teklad Global 18% Protein Rodent Diet (2918, Inotiv, Indianapolis, IN) was studied.

Mice were housed with cellulose bedding (Cellu-nest™, Shepherd Specialty Papers, Watertown, TN) in temperature (∼22°C)- and humidity (30-70%)-controlled conditions maintained on a 14:10 hour light:dark cycle. Both food and water were provided to mice ad libitum. Male mice were used for all experiments and were euthanized via cervical dislocation under one of five conditions: ad libitum fed, 6.5-hour fasted, 18-hour fasted, following a 60-minute treadmill run at 45% of maximal running speed, or following a treadmill run to exhaustion at 22.5 m·min^-1^. Tissues were rapidly excised, freeze-clamped in liquid nitrogen, and stored at -80°C. For 6.5-hour fasting conditions, food was withdrawn during the first hour of the light cycle. For 18-hour fasting conditions, food was removed during the last hour of the light cycle.

### Exercise stress test

The exercise stress test was performed to determine maximal running speed as previously described (7, 16). The maximal running speed obtained is a marker of cardiopulmonary fitness given that it positively correlates with maximum whole body oxygen uptake (16). Mice completed one (for acute exercise experiments) or two (for exercise training experiments) exercise stress tests. For exercise training, the first stress test was completed 24 hours prior to the start of a 6-week exercise training protocol. The second was performed 24 hours following the completion of the 6-week training protocol. Mice were acclimated to running on an enclosed single lane treadmill (Columbus Instruments, Columbus, OH, US) by performing two 10-minute exercise bouts at 10 m·min^-1^ (0% incline) 24 and 48 hours prior to the exercise stress test. For the exercise stress test, mice were placed in the enclosed single lane treadmill. Following a 10-minute sedentary period, mice initiated running at 10 m·min^-1^ (0% incline). The treadmill belt speed was increased by 4 m·min^-1^ every 3 minutes until exhaustion. Exhaustion was defined as the point in which the mouse remained on the shock grid at the back of the treadmill for greater than five seconds.

### Exercise training protocol

The exercise training protocol was completed as previously outlined (8). Briefly, mice began receiving the GAN diet at six weeks of age and the exercise training was initiated when mice were 12 weeks of age. Mice completed a 60-minute treadmill running bout, five days a week for six weeks. The treadmill running speed was 45% of the mouse’s initial maximal running speed during the first two weeks of the training protocol and increased by 5% of the maximal running speed every two weeks. Sedentary mice were placed in an enclosed container on top of the treadmill. Mice were euthanized 72 hours following the final exercise training bout and 48 hours after the final exercise stress test mice.

### Exercise to exhaustion (endurance) test

Mice were acclimated to running on an enclosed single lane treadmill (Columbus Instruments, Columbus, OH, US) as outlined for the exercise stress tests. The endurance test was completed 48 hours after the final acclimation bout. The treadmill belt speed started at 10 m·min^-1^ (0% incline) and increased by ∼4.2 m·min^-1^ every 3 minutes until 22.5 m·min^-1^ was reached. The treadmill speed remained at 22.5 m·min^-1^ until exhaustion.

### Body composition and indirect calorimetry

Body composition was determined in both untrained and trained mice using an EchoMRI-100™ Body Composition Analyzer (EchoMRI LLC, Houston, TX). Indirect calorimetry was performed using the Promethion Core System (Sable Systems International, North Las Vegas, NV). Mice were individually housed for six days on ALPHA-dri® PLUS bedding (Shepherd Specialty Papers, Watertown, TN) in temperature (∼22°C)- and humidity (30-70%)-controlled conditions maintained on a 12:12-hour light:dark cycle. The first three days were the acclimation period followed by three days of data collection to determine food intake, ambulatory activity, voluntary running wheel characteristics, oxygen consumption (VO_2_), CO_2_ production (VCO_2_), respiratory exchange ratio (RER; VCO_2_/VO_2_), and energy expenditure.

Energy expenditure was calculated from VO_2_ and VCO_2_ using the Weir equation (17). Mice used in the indirect calorimetry experiments were 12 weeks of age and had been receiving a GAN diet (D09100310; Research Diets Inc., New Brunswick, NJ) ad libitum for six weeks.

### Surgical procedures

Seventeen-week-old, untrained WT and liver HMGCS2 KO mice had catheters implanted in the jugular vein and carotid artery for isotope infusion and sampling protocols as previously described (8, 18). At 15 weeks of age, a cohort of C57BL/6J mice underwent surgery to implant a carotid artery catheter for measurements of circulating βOHB. The exteriorized ends of the implanted catheters were flushed with 200 U·mL^-1^ heparinized saline and sealed with stainless-steel plugs. Following surgery, mice were housed individually and provided ∼9-10 days of post-operative recovery prior to stable isotope infusion and/or arterial sampling experiments. On post-operative days 6-8, mice underwent acclimation and stress test exercise bouts.

### Stable isotope infusions

Stable isotope infusions were performed at rest and during acute exercise in ∼18-week-old mice as previously described (8). During the first hour of the light cycle, both food and water were withdrawn for the remainder of the experiment (seven hours). Two hours into the fast, mice were moved to an enclosed single lane treadmill and the exteriorized vascular catheters were connected to infusion syringes. Mice were given a one-hour acclimation period prior to obtaining an 80 μl arterial blood sample to determine natural isotopic enrichment of plasma glucose.

Immediately after this sample acquisition, a stable isotope infusion protocol was started to facilitate the quantification of endogenous glucose production and associated oxidative fluxes as previously described (8, 18, 19). Briefly, a ^2^H_2_O (99.9%) bolus containing [6,6-^2^H_2_]glucose (99%) was intravenously infused for 25 minutes to enrich body H_2_O and provide a [6,6-^2^H_2_]glucose prime (440 μmol·kg^-1^). An independent, continuous infusion of [6,6-^2^H_2_]glucose (4.4 μmol·kg^-1^·min^-1^) was initiated after the ^2^H_2_O bolus and [6,6-^2^H_2_]glucose prime. A primed (1.1 mmol·kg^-1^), continuous (0.055 mmol·kg^-1^·min^-1^) intravenous infusion of [U-^13^C]propionate (99%, sodium salt) was started two hours after the ^2^H_2_O bolus and [6,6-^2^H_2_]glucose prime. Four arterial blood samples (100 μl) were obtained 90-120 minutes following the [U-^13^C]propionate bolus (time = 0-30 minutes of treadmill running) to determine arterial glucose, insulin, glucagon, and non-esterified fatty acids (NEFAs). These samples were also used to complete ^2^H/^13^C metabolic flux analysis. The sample acquired at 90 minutes following the [U-^13^C]propionate bolus (time = 0 minutes) was obtained while mice were sedentary on a stationary treadmill.

Samples taken 100-120 minutes following the [U-^13^C]propionate bolus (time = 10-30 minutes of exercise) were acquired while mice were completing an acute treadmill running bout at 45% of their maximal running speed. Plasma samples were stored at -80°C. Donor red blood cells were intravenously infused throughout the experiment to maintain hematocrit. Mice were removed from the treadmill and euthanized by cervical dislocation immediately after the final sample was taken. Liver tissues were rapidly excised, freeze-clamped in liquid nitrogen, and stored at -80°C.

### Glucose derivatization and gas-chromatography-mass spectrometry (GC-MS) analysis

Approximately 40 μl of plasma acquired before starting the stable isotope infusions and at the 0-, 10-, 20-, and 30-minute time points of the treadmill running bout were used to prepare di-*O*-isopropylidene propionate, aldonitrile pentapropionate, and methyloxime pentapropionate derivatives of glucose (7). GC-MS analysis was performed and uncorrected mass isotopomer distributions (MIDs) for six fragment ions were determined as previously described (8, 18).

### 2H/13C metabolic flux analysis

The *in vivo* metabolic flux analysis used in this study has been detailed previously (19, 20). Briefly, a reaction network was generated using Isotopomer Network Compartmental Analysis (INCA) software (21). This reaction network defined both carbon and hydrogen transitions for endogenous glucose production and associated oxidative metabolism reactions. The flux through each network reaction was determined relative to citrate synthase flux (V_CS_) by minimizing the sum of squared residuals between experimentally determined and simulated MIDs of the six fragment ions previously described (20, 22). Flux estimations were repeated 50 times from random initial values. Goodness of fit was assessed by a chi-square test (*p* = 0.0*5*) with 34 degrees of freedom. Body weights of mice and the infusion rate of [6,6-^2^H_2_]glucose were used to determine absolute values.

### Circulating metabolite and hormone analyses

Blood glucose was measured via Contour® blood glucose meters (Ascensia Diabetes Care, Parsippany, NJ). Plasma NEFAs were quantified with the Wako HR series NEFA-HR(2) assay (FUJIFILM Medical Systems USA, Lexington, MA). A Precision Xtra Blood Glucose & Ketone meter (Abbott Diabetes Care, Alameda, CA) was used to measure blood β-ketone bodies. Plasma insulin was determined using the Mercodia Mouse Insulin ELISA (10-1247-01, Winston Salem, NC). Plasma glucagon was quantified via the Mercodia Glucagon ELISA (10-1281-01, Winston Salem, NC).

### Tissue enzyme and metabolite analyses

Citrate synthase activity was assayed at 25 °C as previously described (23). Glycogen was determined by an enzymatic assay as previously described (24). TAGs were determined with the Triglycerides—Liquid Reagent Set (Pointe Scientific, Inc., Lincoln Park, MI). Shotgun lipidomics was performed in liver samples from one cohort of untrained and trained mice to quantify liver TAGs, diacylglycerides (DAGs), cholesterol esters, phosphatidylethanolamine, phosphatidylcholine, and phosphatidylglycerol as previously described (25).

Liver AcAc, βOHB, and total ketone bodies (TKBs; defined as the sum of AcAc and βOHB) were quantified via ultra-high-performance liquid chromatography tandem mass spectrometry (UHPLC-MS/MS) as previously detailed (26). Briefly, liver samples (∼40 mg) were homogenized in an acetonitrile (ACN):methanol (MeOH):water (2:2:1 v/v/v) solution with [U-^13^C_4_]AcAc and D-[3,4,4,4-^2^H_2_] βOHB internal standards. Liver homogenates underwent three cycles of vortexing (10 seconds), flash freezing in liquid nitrogen (30 seconds), and sonication at 25°C (5 minutes). Samples were then centrifuged for 10 minutes at 15,000 x g and 4°C and the supernatants transferred to LC-MS vials. Analysis of samples was performed using a Thermo Fisher Scientific Vanquish LC system and a Cortecs T3 column (186008500, Waters Corporation, Milford, MA) coupled to a Thermo Fisher Scientific QExactive Plus hybrid quadrupole-orbitrap mass spectrometer with a heated electrospray ionization source.

### Immunoblotting

Liver homogenates were prepared as previously described (8, 18, 19). Liver proteins (15 μg) were separated via gel electrophoresis on a NuPAGE 4-12% Bis-Tris gel (Invitrogen, Carlsbad, CA, USA) and transferred to a PVDF membrane. The antibodies used for immunoblotting are provided in Supplemental Table S1. Following incubation with primary and secondary antibodies, PVDF membranes were treated with a chemiluminescent substrate (Thermo Fisher Scientific, Waltham, MA, USA) and images obtained using a ChemiDoc™ Imaging system and Image Lab™ software (Bio-Rad, Hercules, CA, USA). A BLOT-FastStain (G-Bioscience, St. Louis, MO, USA) assay was completed to determine total protein, which was used as the loading control. Densitometry was completed using ImageJ software.

### Statistical analyses

GraphPad Prism software (GraphPad Software LLC., San Diego, CA) was used to perform Student’s t tests, one-way ANOVAs followed by Tukey’s post hoc tests, and two-way ANOVAs and repeated-measures ANOVAs followed by Sidak’s post hoc tests to detect statistical differences (p<0.05). All data are reported as mean ± SEM.

## RESULTS

### Loss of hepatic HMGCS2 promotes liver TAG accumulation during acute exercise

Initial studies evaluated the impact of hepatic HMGCS2 deletion on liver ketone bodies and TAGs across multiple conditions (Fig. 1A-C). Mice were fed a GAN diet for 6 weeks to model the early stages of MASLD. Ketone bodies and TAGs were then quantified in mice that were fed ad libitum, fasted for 6.5 hours, exercised for 60 minutes at 45% of maximal running speed, or exercised to exhaustion. Mice were also fed a GAN diet for 32 weeks to model more advanced stages of MASLD and fasted 6.5 hours. Under each condition, liver HMGCS2 KO mice had lower AcAc, βOHB, and TKBs compared to their WT littermates (Supplemental Fig. S1). When comparing liver ketone bodies within WT mice, mice fasted for 6.5 hours following 6 weeks on a GAN diet showed higher AcAc compared to ad libitum fed mice (Fig. 1 D). Liver AcAc was greater after a 60-minute exercise bout compared to ad libitum fed, exhaustive exercise, and 6.5-hour fasted (32-weeks on GAN diet) conditions (Fig. 1D). Six weeks on a GAN diet followed by a 6.5-hour fast elevated βOHB and TKBs versus ad libitum fed mice and mice fasted 6.5 hours following 32 weeks on a GAN diet (Fig. 1E and F). Sixty minutes of treadmill running increased liver βOHB and TKBs relative to all other conditions in WT mice (Fig. 1E and F). Liver βOHB in WT mice was higher following exhaustive exercise compared to the fed state (Fig. 1E).

**Figure 1.**
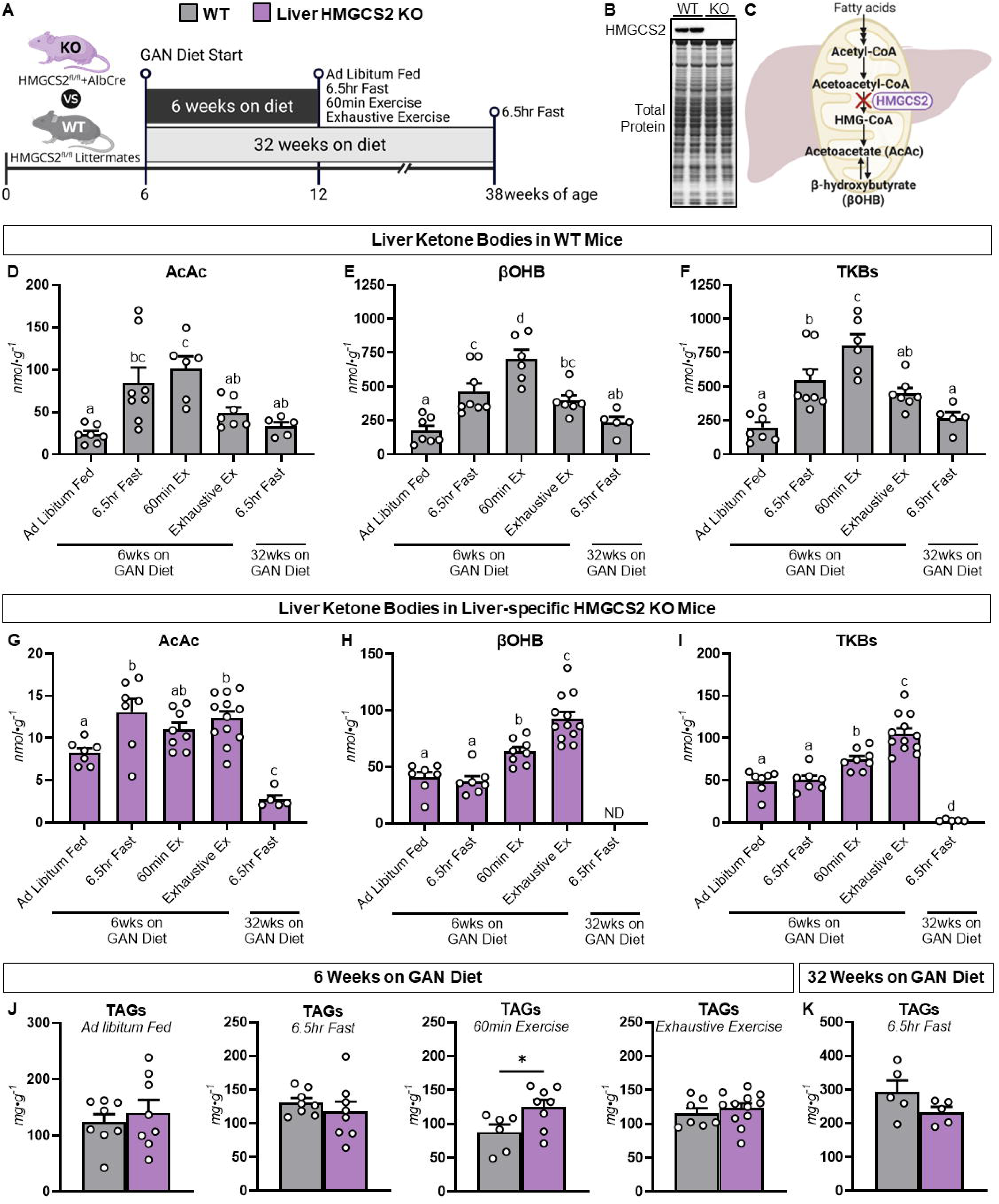
Loss of HMGCS2 promotes liver triacylglyceride accumulation during acute exercise. **A:** A schematic representation of the experimental design. Liver-specific HMGCS2 knockout (KO) mice and wild type (WT) littermates were fed a GAN diet for 6 or 32 weeks. Tissues were collected from mice that were fed ad libitum, fasted for 6.5 hours, exercised on a treadmill for 60 minutes at 45% of their maximal running speed, or exercised on a treadmill at 22.5 m·min^-1^ until exhaustion. **B:** A representative immunoblot of liver HMGCS2 in WT and liver-specific HMGCS2 KO mice. **C:** A schematic representation the ketogenic pathway with select metabolites and enzymes. Liver acetoacetate (AcAc; nmol·g^-1^), β-hydroxybutyrate (βOHB; nmol·g^-1^), and total ketone bodies (TKBs; nmol·g^-1^) in WT mice (**D-F**; n = 5-8 per genotype). Liver AcAc (nmol·g^-1^), βOHB (nmol·g^-1^), and TKB (nmol·g^-1^) in liver HMGCS2 KO mice (**G-I**; n = 5-12). **J:** Liver triacylglycerides (TAGs; mg·g^-1^) in mice fed a GAN diet for 6 weeks (n = 6-12 per genotype). **K:** Liver TAGs (mg·g^-1^) in mice fed a GAN diet for 32 weeks (n = 5 per genotype). Data are mean ± SEM. Letters denote significant differences between groups (p<0.05). *p<0.05 between specified groups. ND = not determined because concentrations are below quantifiable limits.

Within KO mice, liver AcAc was higher in mice fasted 6.5 hours following 6 weeks on a GAN diet compared to ad libitum fed mice and mice fasted 6.5 hours following 32 weeks on a GAN diet (Fig. 1G). Sixty minutes of treadmill running increased liver AcAc relative to 6.5-hour fasted mice following 32 weeks on a GAN diet (Fig. 1G). Mice completing exhaustive exercise showed elevated AcAc compared to ad libitum fed mice and mice fasted 6.5 hours following 32 weeks on diet (Fig. 1G). Liver βOHB and TKBs were increased in mice following 60 minutes of treadmill running compared to ad libitum fed and 6.5-hour fasted mice (Fig. 1H and I). Exhaustive exercise increased liver βOHB and TKBs relative to all groups (Fig. 1H and I).

Notably, only the mice performing a 60-minute exercise bout showed a genotype effect for liver TAGs. Specifically, liver TAGs were higher in liver HMGCS2 KO mice compared to WT littermates (Fig. 1J and K). Together, our results show that the increase in liver ketone bodies by submaximal exercise (i.e., 60-minute exercise group) is impeded by loss of hepatic HMGCS2 and accompanied by elevated liver TAGs. Our results also support that exercise (60-minute and exhaustive exercise) promotes a rise in liver ketone bodies from both HMGCS2-dependent and -independent sources and long-term high-fat feeding (32-week GAN diet) lowers liver ketone bodies, in part, by decreasing those sourced independently of HMGCS2.

### Hepatic HMGCS2 KO promotes liver TAG accumulation under conditions of acutely elevated lipid availability

Submaximal exercise is accompanied by increased lipid delivery to the liver, which could place greater reliance on hepatic ketogenesis to maintain liver lipid homeostasis. To further test if hepatic HMGCS2 was required for liver lipid homeostasis during short-term elevations in liver lipid availability, WT and liver HMGCS2 KO mice were fed a GAN diet for 6 weeks and fasted for 18 hours. Body weight was comparable between genotypes (Fig. 2A). Liver AcAc, βOHB, and TKBs were decreased in liver HMGCS2 KO mice compared to WT mice (Fig. 2B). Moreover, mice lacking hepatic HMGCS2 had elevated liver TAGs relative to WT littermates (Fig. 2C).

**Figure 2.**
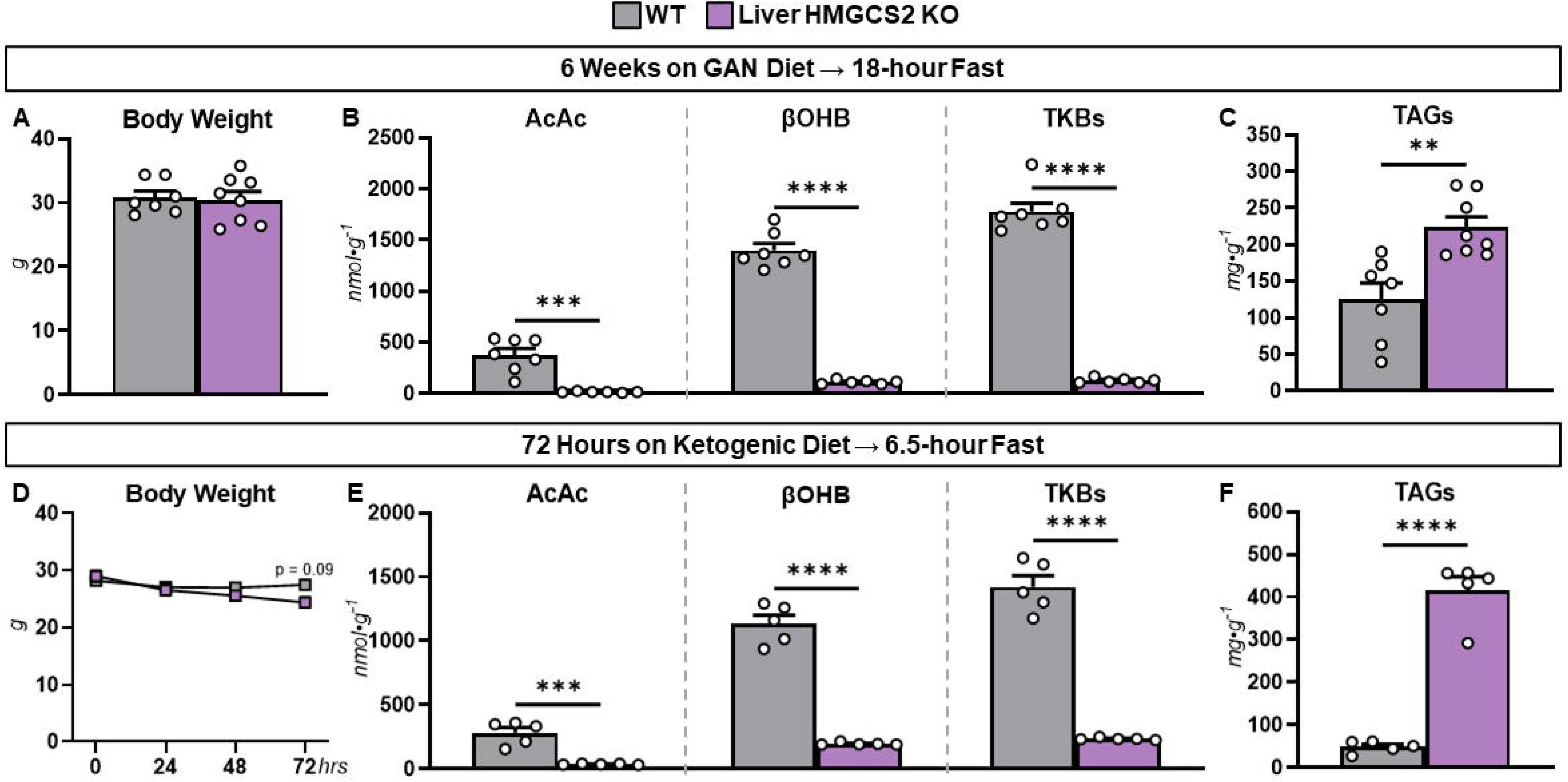
Loss of HMGCS2 promotes liver triacylglyceride accumulation under conditions of acutely elevated lipid availability. **A:** Body weights of 12-week-old, liver-specific HMGCS2 knockout (KO) mice and wild type (WT) littermates fed a GAN diet for 6 weeks (n = 7-8 per genotype). **B:** Liver acetoacetate (AcAc; nmol·g^-1^), β-hydroxybutyrate (βOHB; nmol·g^-1^), and total ketone bodies (TKBs; nmol·g^-1^) in 18-hour fasted WT and liver-specific HMGCS2 KO mice following 6 weeks on a GAN diet (n = 6-7 per genotype). **C:** Liver triacylglycerides (TAGs; mg·g^-1^) in 18-hour fasted WT and liver-specific HMGCS2 KO mice following 6 weeks on a GAN diet (n = 7-8 per genotype). **D:** Body weights of 12-week-old, liver-specific HMGCS2 KO mice and WT littermates fed a ketogenic diet for 72 hours (n = 5 per genotype). **E:** Liver AcAc (nmol·g^-1^), βOHB (nmol·g^-1^), and TKBs (nmol·g^-1^) in 6.5-hour fasted WT and liver-specific HMGCS2 KO mice following 72 hours on a ketogenic diet (n = 5 per genotype). **F:** Liver TAGs (mg·g^-1^) in 6.5-hour fasted WT and liver-specific HMGCS2 KO mice following 72 hours on a ketogenic diet (n = 5 per genotype). Data are mean ± SEM. **p<0.01, ***p<0.001, and ****p<0.0001.

We also fed WT and liver HMGCS2 KO mice a ketogenic diet (90.5% kcal from fat) for 72 hours to acutely increase liver lipid availability from an exogenous source. Body weight was similar between genotypes over the 72 hours on the ketogenic diet (Fig. 2D). As expected, liver AcAc, βOHB, and TKBs were lower in KO mice compared to WT controls (Fig. 2E). Liver TAGs were ∼7.75-fold higher in liver HMGCS2 KO mice relative to WT mice (Fig. 2F). These results are consistent with the results from the acute exercise experiments (Fig. 1). That is, under conditions characterized by short-term elevations in lipid availability, hepatic HMGCS2 is necessary to prevent liver TAG accumulation.

### Loss of hepatic BDH1 does not promote liver TAG accumulation during acute exercise

Mice lacking liver BDH1 were studied to define the importance of disposing of fatty acids to AcAc on liver lipid homeostasis during exercise (Fig. 3A). Body weight, fat mass, and lean mass were similar between genotypes (Fig. 3B). Loss of hepatic BDH1 did not impact maximal running speed (Fig. 3C). Following a 60-minute treadmill run, liver AcAc was higher, βOHB was decreased, and TKBs were unchanged in liver BDH1 KO compared to WT mice (Fig. 3D). Notably, liver TAGs were similar between liver BDH1 KO mice and WT littermates (Fig. 3E). To further support our findings from the acute exercise experiments, WT and liver BDH1 KO mice were fasted 18 hours. Loss of hepatic BDH1 increased AcAc, decreased βOHB, and lowered TKBs in 18-hour fasted mice (Fig. 3F). Liver TAGs were comparable between WT and liver BDH1 KO mice (Fig. 3G). These results suggest that the ability to divert acetyl-CoA and/or dispose of fatty acids towards AcAc is important for maintaining liver lipid homeostasis during acute exercise.

**Figure 3.**
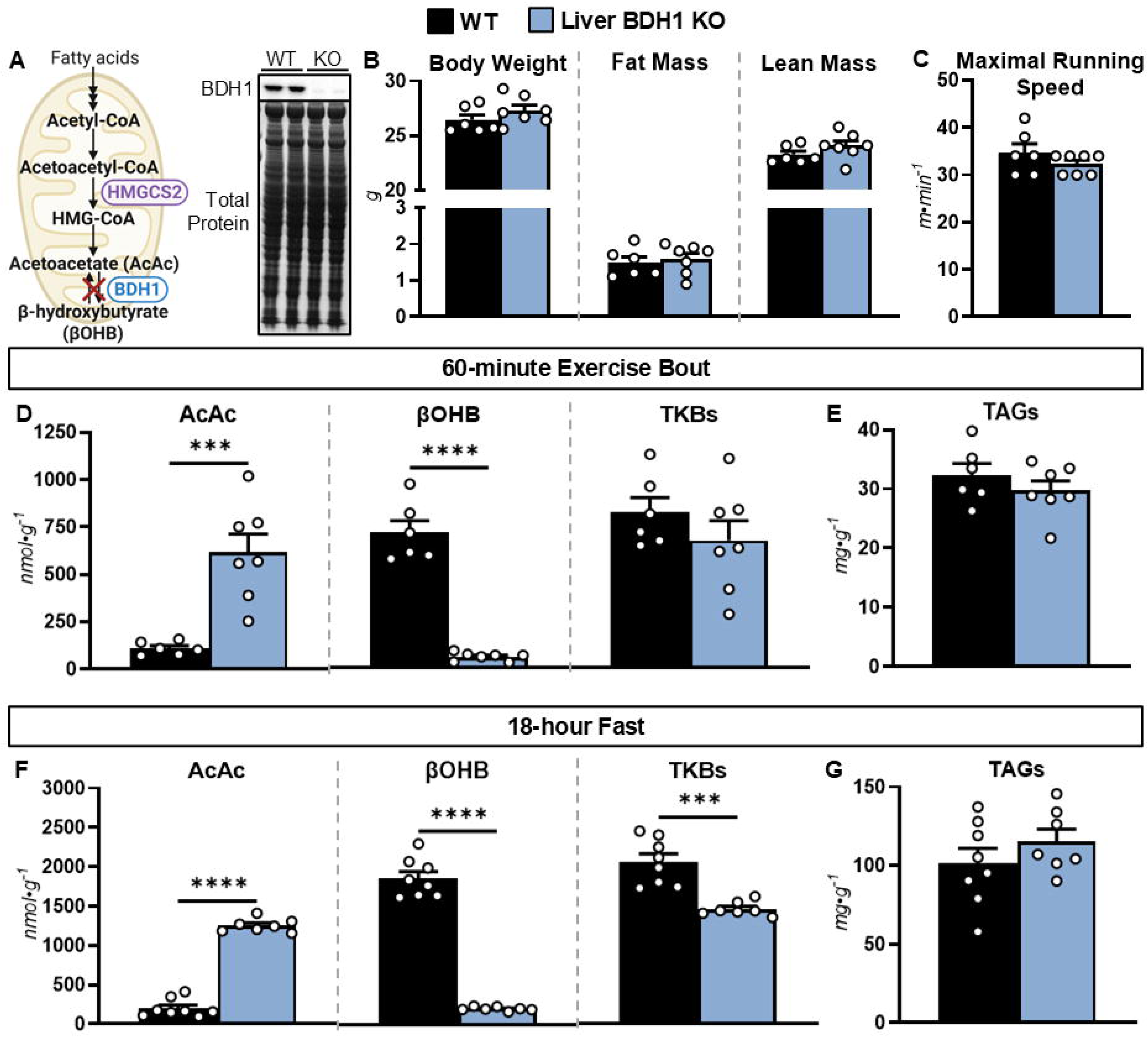
Liver BDH1 knockout does not increase liver triacylglycerides under conditions of acutely elevated lipid availability. **A:** A schematic representation the ketogenic pathway with select metabolites and enzymes and a representative immunoblot of liver BDH1 in liver-specific BDH1 knockout (KO) mice and wild type (WT) littermates. **B:** Body weight (g), fat mass (g), and lean mass (g) in 12-week-old, chow-fed WT and liver BDH1 KO mice (n = 6-7 per genotype). **C:** Maximal running speed in (m·min^-1^) in WT and liver BDH1 KO mice (n = 6-7 per genotype). **D:** Liver acetoacetate (AcAc; nmol·g^-1^), β-hydroxybutyrate (βOHB; nmol·g^-1^), and total ketone bodies (TKBs; nmol·g^-1^) in WT and liver-specific BDH1 KO mice following a 60-minute treadmill run at 45% of maximal running speed (n = 6-7 per genotype). **E:** Liver triacylglycerides (TAGs; mg·g^-1^) in WT and liver-specific BDH1 KO mice following a 60-minute treadmill run at 45% of maximal running speed (n = 6-7 per genotype). **F:** Liver AcAc (nmol·g^-1^), βOHB (nmol·g^-1^), and TKBs (nmol·g^-1^) in 18-hour fasted WT and liver-specific BDH1 KO mice (n = 7-8 per genotype). **G:** Liver TAGs (mg·g^-1^) in 18-hour fasted WT and liver-specific BDH1 KO mice (n = 7-8 per genotype). Data are mean ± SEM. ***p<0.001, and ****p<0.0001.

### TCA cycle flux is higher in hepatic HMGCS2 KO mice at rest, but not with increasing exercise duration

In addition to disposal by ketogenesis, acetyl-CoA and fatty acids are also oxidized by the TCA cycle (in conjunction with oxidative phosphorylation) to support the energetic demands of gluconeogenesis. As such, TCA cycle fluxes were quantified in mice at rest and during a 30-minute exercise bout (Fig. 4A) to further test the impact of ketogenic insufficiency on lipid homeostasis. Plasma insulin decreased similarly in WT and liver HMGCS2 KO mice during exercise (Fig. 4B). Plasma glucagon increased comparably in both genotypes during the 30-minute treadmill run (Fig. 4C). Plasma NEFAs were higher in liver HMGCS2 KO mice at rest, but decreased during exercise to concentrations comparable to that of WT mice (Fig. 4D). Arterial glucose was similar between liver HMGCS2 KO mice and WT littermates at rest and throughout the exercise bout (Fig. 4E).

**Figure 4.**
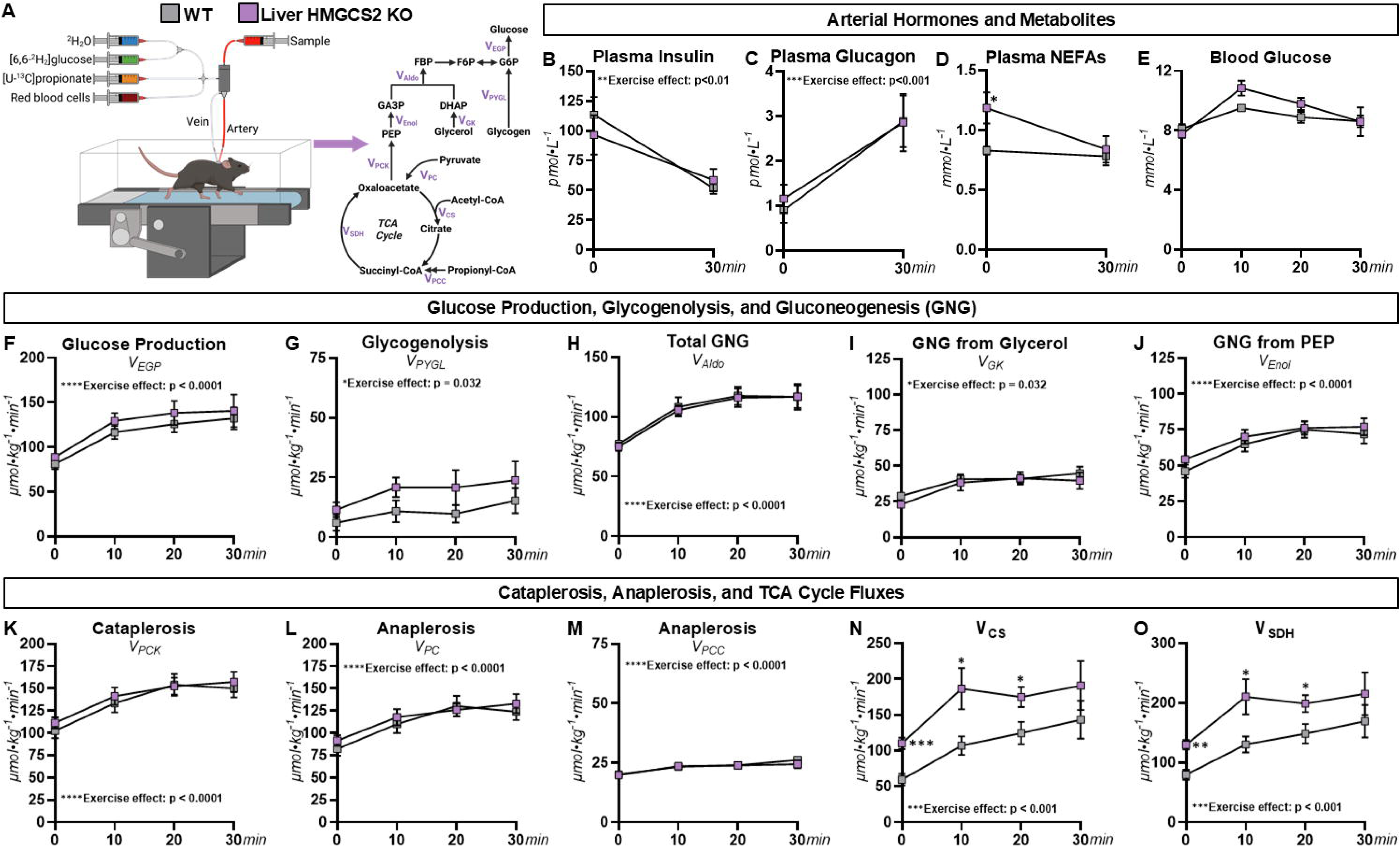
Glucose and oxidative fluxes at rest and during acute exercise in mice lacking liver HMGCS2. Eighteen-week-old, liver-specific HMGCS2 knockout mice and wild type (WT) littermates fed a GAN diet for 12 weeks underwent ^2^H- and ^13^C-isotope infusions to quantify glucose and oxidative fluxes at rest and during an acute exercise bout **(A)**. **B:** Plasma insulin (pmol·L^-1^) prior to and at the conclusion of a 30-minute treadmill run (n = 6-7 per genotype). **C:** Plasma glucagon (pmol·L^-1^) prior to and at the conclusion of a 30-minute treadmill run (n = 7 per genotype). **D:** Plasma non-esterified fatty acids (NEFAs; mmol·L^-1^) prior to and at the conclusion of a 30-minute treadmill run (n = 7 per genotype). **E:** Blood glucose (mmol·L^-1^) prior to and during a 30-minute treadmill run (n = 6-7 per genotype). Model-estimated, absolute nutrient fluxes (μmol·kg^-1^·min^-1^) in WT and liver-specific HMGCS2 KO mice (n = 6-7 per genotype) prior to and during a 30-minute of treadmill run for **(F)** endogenous glucose production (V_EGP_), **(G)** glycogenolysis (V_PYGL_), **(H)** total gluconeogenesis (V_Aldo_), **(I)** gluconeogenesis from glycerol (V_GK_), **(J)** gluconeogenesis from phosphoenolpyruvate (V_Enol_), **(K)** tricarboxylic acid cycle cataplerosis (V_PCK_), **(L)** anaplerosis from pyruvate (V_PC_), **(M)** anaplerosis from propionyl-CoA (V_PCC_), **(N)** flux from oxaloacetate and acetyl-CoA to citrate (V_CS_), and **(O)** flux from succinyl-CoA to oxaloacetate (V_SDH_). Data are mean ± SEM. *p<0.05, **p<0.01, and ***p<0.001 vs. WT mice at the specified time point.

Endogenous glucose production (V_EGP_), glycogenolysis (V_PYGL_), and total gluconeogenesis (V_Aldo_) were similar at rest and increased during exercise in both genotypes (Fig. 4F-H). The rise in total gluconeogenesis during muscular work in WT and liver HMGCS2 KO mice was due to elevated rates of gluconeogenesis from glycerol (V_GK_), gluconeogenesis from phosphoenolpyruvate (V_Enol_), and cataplerosis from the TCA cycle (Fig. 4I-K). Anaplerosis was also higher in response to exercise in both WT and liver HMGCS2 KO mice (Fig. 4L and M). This included flux of pyruvate to oxaloacetate (V_PC_; Fig. 4L) and flux from propionyl-CoA to succinyl-CoA (V_PCC_; Fig. 4M). TCA cycle fluxes (V_CS_ and V_SDH_) were elevated in liver HMGCS2 KO mice compared to WT littermates at rest (Fig. 4N and O). TCA cycle fluxes increased during muscular work in both genotypes (Fig. 4N and O). However, V_CS_ and V_SDH_ were no longer higher in KO mice relative to WT mice at 30 minutes of treadmill running (Fig. 4 N and O). These data indicate that enhanced hepatic disposal of lipids via the TCA cycle may compensate for ketogenic insufficiency and prevent liver TAG accumulation in liver HMGCS2 KO mice under sedentary conditions, but not during exercise.

### Loss of hepatic HMGCS2 increases voluntary wheel running

Next, we performed indirect calorimetry to assess the extent to which dysregulated liver oxidative metabolism in liver HMGCS2 KO mice impacted systemic energy metabolism. Oxygen consumption (VO_2_; Fig. 5A), CO_2_ production (VCO_2_; Fig. 5B), RER (Fig. 5C) and energy expenditure (Fig. 5D) were similar between liver HMGCS2 KO mice and WT controls during both the light and dark cycles. VO_2_, VCO_2_, and energy expenditure were higher in both genotypes during the dark cycle compared to the light cycle (Fig. 5A, B, and D). In contrast, RER increased in the dark cycle compared to the light cycle in liver HMGCS2 KO mice but not WT mice (Fig. 5D). Body weight was not different between genotypes (Fig. 5E). Food intake was comparable between WT and liver HMGCS2 KO mice during both photoperiods (Fig. 5F). Ambulatory movement off the voluntary running wheel was similar between WT and KO mice (Fig. 5G). Voluntary running wheel distance was also comparable between genotypes during the light cycle (Fig. 5H). In contrast, liver HMGCS2 KO mice displayed a higher voluntary running wheel distance during the dark cycle (Fig. 5H). This increased wheel running distance in mice lacking hepatic HMGCS2 was associated with a greater amount of time spent running at wheels speeds above 25 m·min^-1^ in the dark cycle (Fig. 5I and J). To further test if the differences in wheel running characteristics between genotypes were voluntary or physiological in nature, mice underwent a forced treadmill run to exhaustion. Time to exhaustion running at 22.5 m·min^-1^ was similar in WT and liver HMGCS2 KO mice (Fig. 5K). These results suggest the increased voluntary wheel running distance and speeds in liver HMGCS2 KO mice are not owing to heightened exercise endurance.

**Figure 5.**
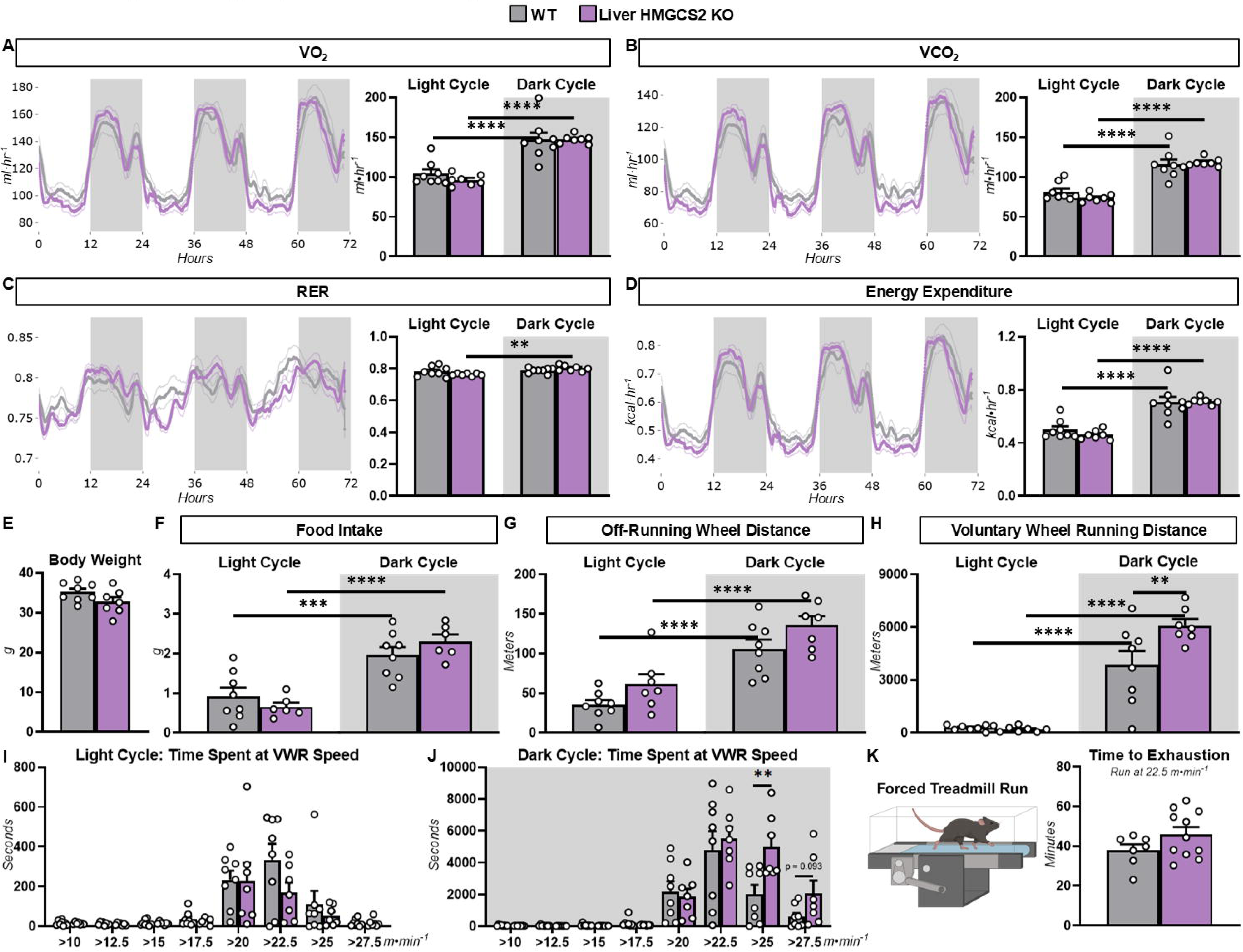
Voluntary wheel running (VWR) is increased in mice lacking liver HMGCS2. Twelve-week-old liver specific HMGCS2 knockout mice and wild type (WT) littermates were fed a GAN diet for 6 weeks prior to indirect calorimetry and voluntary wheel running experiments. **A:** Oxygen consumption (VO_2_; ml·hr^-1^; n = 7-8 per genotype). **B:** Carbon dioxide production (VCO_2_; ml·hr^-1^; n = 7-8 per genotype). **C:** Respiratory exchange ratio (RER; unitless; n = 7-8 per genotype). **D:** Energy expenditure (kcal·hr^-1^; n = 7-8 per genotype). **E:** Body weight (g; n = 7-8 per genotype). **F:** Food intake (g; n = 6-8 per genotype). **G:** All meters by mice while not on the voluntary running wheel (n = 7-8 per genotype). **H:** Running distance (meters) while on the voluntary running wheel (n = 7-8 per genotype). **I:** Time spent (seconds) at different voluntary wheel running speeds during the light cycle (n = 7-8 per genotype). **J:** Time spent (seconds) at different voluntary wheel running speeds during the dark cycle (n = 7-8 per genotype). **K:** Time to exhaustion (minutes) while running on a treadmill at 22.5 m·min^-1^ (n = 7-11 per genotype). Data are mean ± SEM. **p<0.01, ***p<0.001, and ****p<0.0001.

### Exercise training limits gains in body weight and adiposity in both WT and KO mice

Given that ketogenic insufficiency dysregulated liver lipid homeostasis during acute exercise, the impact of exercise training on systemic and tissue-specific lipid metabolism was tested in WT and liver HMGCS2 KO mice. The exercise training protocol was 60 minutes of treadmill running, five days per week for six weeks (Fig. 6A). The treadmill running speed was 45% of the mouse’s initial maximal running speed during the first two weeks of the training protocol and increased by 5% of the maximal running speed every two weeks. This initial speed and duration were selected because they stimulate an increase in circulating βOHB during exercise that persists upon completion of the exercise bout (Fig. 6B). Ad libitum body weight, adiposity, and lean mass were comparable between WT and KO mice prior to exercise training protocols at 12 weeks of age (Fig. 6C). Reassessment of these metrics following 6 weeks of exercise training showed that trained mice in both genotypes had lower body weight, fat mass and lean mass compared to their untrained counterparts (Fig. 6D). While six weeks of exercise training prevented a gain in body weight and adiposity in both genotypes, the protective effect of training was delayed in liver HMGCS2 KO mice (Fig. 6E and F).

**Figure 6.**
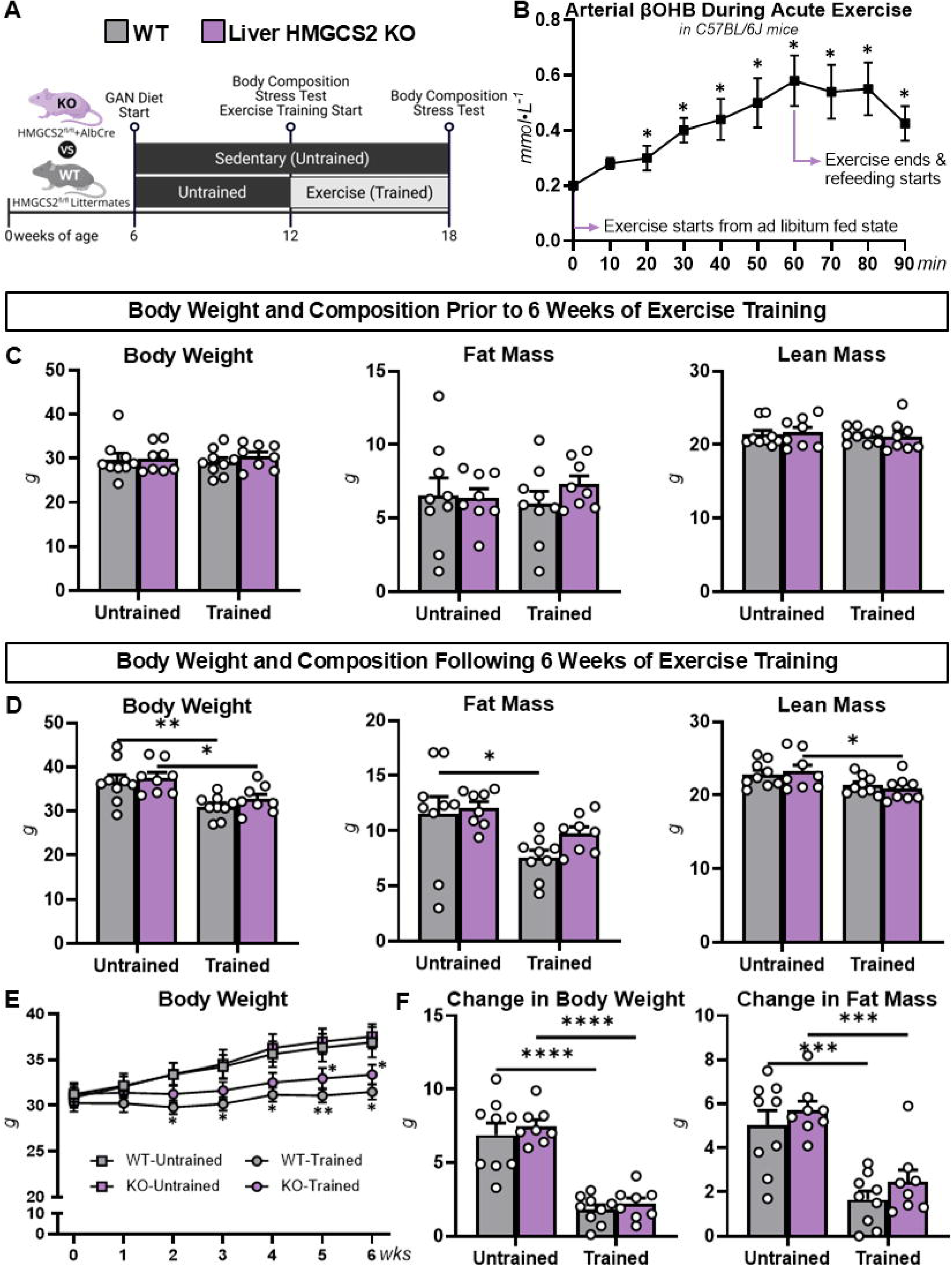
Anthropometric adaptations to exercise training in mice lacking liver HMGCS2. **A:** A schematic representation of the exercise training protocol. Liver-specific HMGCS2 knockout (KO) mice and wild type (WT) littermates were fed a GAN diet starting at 6 weeks of age. At 12 weeks of age, mice were randomly assigned to sedentary (untrained) or exercise training (trained) groups. **B:** Arterial blood β-hydroxybutyrate (βOHB; mmol·L^-1^) in C57BL/6J mice during and following a treadmill run at 45% of maximal running speed (n = 4-5 mice per time point; *p<0.05 vs. 0-minute time point). **C:** Body weight (g), fat mass (g), and lean mass (g) prior to the exercise training protocol (n = 8-9 per genotype). **D:** Body weight (g), fat mass (g), and lean mass (g) following the exercise training protocol (n = 8-9 per genotype). **E:** A time course of body weight (g) during the 6-week exercise training protocol (n = 8-9 per genotype). **F:** Change in body weight (g) and fat mass (g) during the 6-week exercise training protocol (n = 8-9 per genotype). Data are mean ± SEM. **C-F:** *p<0.05, **p<0.01, ***p<0.001, and ****p<0.0001 vs. untrained mice within genotypes.

### Lowering of liver fat by exercise training is not compromised in mice lacking hepatic HMGCS2

Experiments also tested the impact of exercise training on liver metabolism in WT and liver HMGCS2 KO mice (Fig. 7). Liver HMGCS2 protein was comparable in untrained and trained WT mice (Fig. 7A). Liver BDH1 protein was similar in untrained WT and KO mice (Fig. 7A). Training decreased liver BDH1 in both genotypes (Fig. 7A). As expected, liver AcAc, βOHB, and TKBs were lower in untrained and trained KO mice compared to their WT controls (Fig. 7B). Trained WT mice had decreased liver AcAc, βOHB, and TKBs relative to untrained WT mice (Fig. 7B). Liver TAGs were similar in untrained WT and KO mice (Fig. 7C). Training decreased liver TAGs in both genotypes (Fig. 7C). In addition to total concentration, we assessed the fatty acyl chains comprising liver TAGs. Training decreased the majority of fatty acids quantified in both WT and liver HMGCS2 KO mice (Fig. 7C). Liver DAGs were similar in untrained WT and KO mice (Fig. 7D). Training diminished DAGs in WT, but not liver HMGCS2 KO mice (Fig. 7D). Liver cholesterol esters (Fig. 7E), phosphatidylethanolamine (Fig. 7F), phosphatidylcholine (Fig. 7G), and phosphatidylglycerol (Fig. 7H) were comparable between all groups. The key finding related to liver lipid metabolism presented here is that hepatic HMGCS2 is not necessary for exercise training to lower liver TAGs.

**Figure 7.**
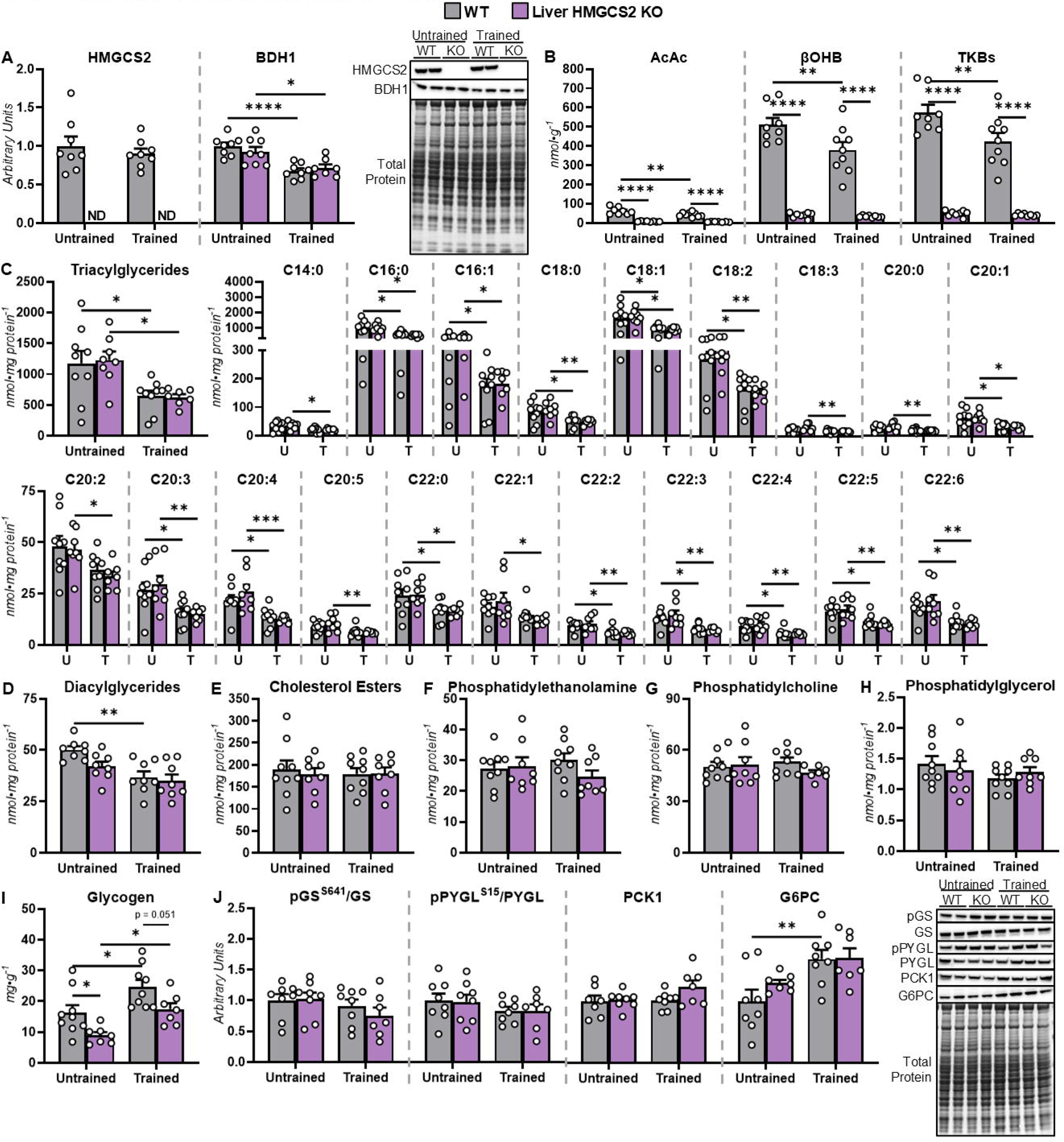
Liver adaptations to exercise training in mice lacking liver HMGCS2. Livers were analyzed in 18-week-old untrained (U) and exercise-trained (T) mice with a liver-specific HMGCS2 knockout (KO) and wild type (WT) littermates fed a GAN diet starting at 6 weeks of age. **A:** Liver HMGCS2 and BDH1 as determined by immunoblotting and representative immunoblots (arbitrary units; n = 7-8 per genotype). **B:** Liver acetoacetate (AcAc; nmol·g^-1^), β-hydroxybutyrate (βOHB; nmol·g^-1^), and total ketone bodies (TKBs; nmol·g^-1^). **C:** Liver triacylglycerides and fatty acids comprising triacylglycerides (nmol·mg protein^-1^; n = 7-9 per genotype). **D:** Liver diacylglycerides (nmol·mg protein^-1^; n = 8 per genotype). **E:** Liver cholesterol esters (nmol·mg protein^-1^; n = 8-9 per genotype). **F:** Liver phosphatidylethanolamine (nmol·mg protein^-1^; n = 8-9 per genotype). **G:** Liver phosphatidylcholine (nmol·mg protein^-1^; n = 7-9 per genotype). **H:** Liver phosphatidylglycerol (nmol·mg protein^-1^; n = 8-9 per genotype). **I:** Liver glycogen (nmol·mg protein^-1^; n = 7-9 per genotype). **J:** Liver phospho-glycogen synthase (pGS)-to-total GS ratio, phospho-glycogen phosphorylase (pPYGL)-to-total PYGL ratio, cytosolic phosphoenolpyruvate carboxykinase (PCK1), and glucose-6-phosphatase catalytic subunit (G6PC) as determined by immunoblotting and representative immunoblots (arbitrary units; n = 7-8 per genotype). Data are mean ± SEM. *p<0.05, **p<0.01, ***p<0.001, and ****p<0.0001

In addition to lipid characteristics, indices of liver glycogen metabolism were evaluated in untrained and trained mice. Liver glycogen was lower in untrained KO mice compared to untrained WT mice (Fig. 7I). Training elevated liver glycogen in both WT and liver HMGCS2 KO mice (Fig. 7I). However, glycogen in livers of trained WT mice was greater than that of trained KO mice (Fig. 7I). Molecular regulators of glycogen metabolism were assessed. The liver phospho-glycogen phosphorylase (pPYGL)-to-PYGL ratio and phospho-glycogen synthase (pGS)-to-GS ratio did not differ between genotypes or in response to training (Fig. 7J).

Regulators of glucose-6-phosphate, a glycogen precursor/product, were determined. Phosphoenolpyruvate carboxykinase 1 (PCK1) protein was unaffected (Fig. 7J). A training effect showed liver glucose-6-phosphatase (G6PC) protein was higher in trained WT mice compared to untrained WT mice (Fig. 7J). These results indicate ketogenic insufficiency lowers liver glycogen, but does not impede the training-induced increase in liver glycogen deposition.

### Skeletal muscle adaptations to exercise training may be blunted in mice lacking liver HMGCS2

Given that liver-sourced ketone bodies can support the energetic demands of muscular work, adaptations in exercise performance and metabolism were assessed in WT and liver HMGCS2 KO mice following six weeks of exercise training. Maximal running speed and total work during an exercise stress test were similar across all groups prior to exercise training (Fig. 8A). In contrast, training significantly improved maximal running speed in WT, but not KO, mice (Fig. 8B). Citrate synthase activity (CSA) in the gastrocnemius was similar between untrained WT and liver HMGCS2 KO mice (Fig. 8C). Training elevated gastrocnemius CSA in WT mice, but did not significantly increase CSA in KO mice (Fig. 8C). Gastrocnemius glycogen was comparable between all groups (Fig. 8D). Gastrocnemius TAGs were not different between untrained WT and liver HMGCS2 KO mice (Fig. 8E). Trained WT mice had lower gastrocnemius TAGs relative to untrained WT mice (Fig. 8E). This training effect on gastrocnemius TAGs was not significant in liver HMGCS2 KO mice (Fig. 8E). Superficial vastus lateralis (SVL) TAGs were similar between all groups (Fig. 8F). However, SVL TAGs trended higher in trained KO compared to trained WT mice (Fig. 8F). Together these data suggest ketogenic insufficiency may have a negative, albeit modest, impact on muscle adaptations to exercise training.

**Figure 8.**
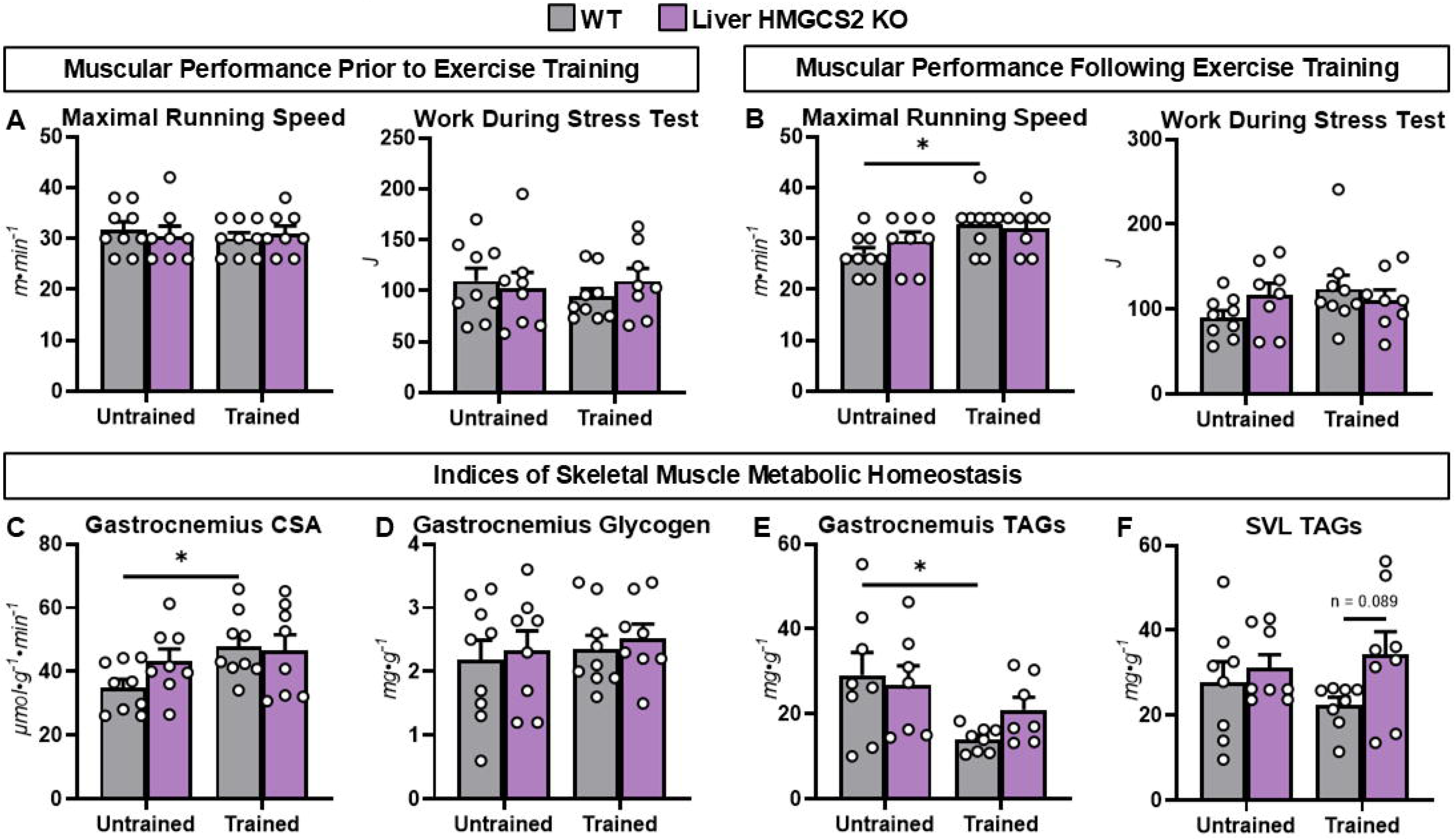
Skeletal muscle adaptations to exercise training in mice lacking liver HMGCS2. **A:** Maximal running speed (m·min^-1^) and work (J) during an exercise stress test in 12-week-old mice with a liver-specific knockout (KO) of HMGCS2 and wild type (WT littermates prior to 6 weeks of exercise training protocols (n = 8-9 per group). **B:** Maximal running speed (m·min^-1^) and work (J) during an exercise stress test in 18-week-old mice following 6 weeks of exercise training protocols (n = 8-9 per group). Skeletal muscle was analyzed in 18-week-old untrained and exercise-trained mice. **C:** Gastrocnemius citrate synthase activity (CSA; µmol·g^-1^·min^-1^; n = 8-9 per genotype). **D:** Gastrocnemius glycogen (mg·g^-1^; n = 8-9 per genotype). **E:** Gastrocnemius triacylglycerides (TAGs; mg·g^-1^; n = 7-8 per genotype). **F:** Superficial vastus lateralis triacylglycerides (TAGs; mg·g^-1^; n = 8 per genotype). Data are mean ± SEM. *p<0.05.

## DISCUSSION

Acute exercise stimulates hepatic lipid disposal pathways that transform the potential energy in fatty acids to alternative forms of energy currency that are then released from the liver to support muscular work. This includes increasing the production of AcAc and βOHB from fatty acids via ketogenesis (11). A gap in knowledge has been whether the provocation of hepatic ketogenesis by acute exercise is critical for exercise training to mitigate liver steatosis. In this study we tested the hypothesis that hepatic ketogenesis is necessary for exercise training to lower liver lipids. The key findings of our work are i) loss of hepatic HMGCS2 promotes liver lipid accumulation during acute, non-exhaustive exercise and ii) hepatic HMGCS2-dependent ketogenesis is not necessary for exercise training to lower liver lipids.

### Role of ketogenic insufficiency in liver lipid homeostasis during acute exercise

Exercise stimulates ketogenesis through i) the mobilization of fatty acids from adipose tissue and delivery to the liver, ii) the hepatic extraction of fatty acids from the circulation, and iii) the intrahepatic conversion of fatty acids to ketone bodies (4). Studies in both humans and experimental models support that the fall in insulin during exercise promotes all three of these processes (4, 27–29). In addition, the exercise-mediated rise in glucagon enhances the conversion of fatty acids to both AcAc and βOHB within the liver (4, 13). Consistent with this prior work, our data show that exercise lowers insulin, increases glucagon, and elevates liver ketone bodies in WT mice. Liver HMGCS2 KO mice exhibited a similar hormonal response to exercise compared to WT mice. However, impeding the conversion of fatty acids to ketone bodies via deletion of hepatic HMGCS2 led to a blunted rise in liver ketone bodies and increased liver TAGs in response to non-exhaustive exercise. To further test the role of ketogenic insufficiency in promoting liver lipid accumulation during exercise, we studied hepatic BDH1 KO mice to permit the generation of AcAc, but not βOHB, from fatty acids. In contrast to hepatic HMGCS2 KO mice, loss of BDH1 in livers of mice did not result in an exercise-mediated rise in liver TAGs. Our results indicate that HMGCS2 is a key node through which ketogenesis promotes liver lipid homeostasis during non-exhaustive exercise.

While the deletion of hepatic HMGCS2 inhibits the production of ketone bodies from fatty acids within the liver, ketogenic insufficiency is unlikely to be solely responsible for the accumulation of liver fat in HMGCS2 KO mice during exercise. Here we show that, under sedentary conditions, ad libitum fed and 6.5-hour fasted mice lacking hepatic HMGCS2 had lower liver ketone bodies but similar liver TAG concentrations compared to WT controls. Prior work using adult mice with a whole-body and liver-specific HMGCS2 KO or knockdown have also reported that ketogenic insufficiency does not promote liver steatosis in fed and short-fasted (<8 hours) states (14, 30–33). These results indicate that the inhibition of hepatic ketogenesis alone is not sufficient to promote liver steatosis. As previously mentioned, hepatic fatty acid delivery and ketone body production are positively correlated during exercise (34). The current study had mice undergo an 18-hour fast and short-term ketogenic diet feeding because they lower circulating insulin and acutely increase liver lipid availability (35–39). Under these conditions, liver HMGCS2 KO mice had increased liver TAGs relative to WT mice. Our results are in agreement with independent studies demonstrating that long fasting durations (>8 hours) and very high fat diets (>90.5%) elevate liver TAGs in mice lacking HMGCS2 (14, 31). The interaction between ketogenic insufficiency and lipid availability on liver steatosis can also be gleaned from studies in neonatal mice. Prior to postnatal day 14, mice with a whole-body knockdown or KO of HMGCS2 display fatty liver (33, 40). This may be due to the primary nutrient source for rodents during this period of life being breast milk, which is high in fat content (40). Notably, this liver steatosis phenotype in ketogenesis-insufficient neonatal mice is lost when transitioned to a high-carbohydrate, low-fat diet (33). It is currently unclear why the GAN diet used in our study did not promote fatty liver in sedentary HMGCS2 KO mice. While the GAN diet increases lipid availability (40% kcal from fat), it also elevates circulating insulin levels in mice (41). Wasserman et al. identified that preventing the fall in insulin during exercise attenuates hepatic extraction of fatty acids and stimulation of ketogenesis (27). We showed that liver-specific HMGCS2 KO mice had elevated circulating NEFAs at rest (when insulin is high), but not during exercise (when insulin is low). It may be the higher insulin levels in mice fed a GAN diet limit hepatic extraction of lipids in sedentary, liver HMGCS2 KO mice. Together, our data and the work of others suggest that loss of hepatic HMGCS2 provokes liver lipid accumulation during acute exercise because the inability to convert fatty acids to ketone bodies in the liver is accompanied by a concurrent increase in lipid delivery to the liver and/or hepatic extraction of lipids from the circulation.

Conditional compensation by other lipid disposal pathways could contribute to our observations that mice lacking hepatic HMGCS2 are protected from fatty liver at rest, but not during exercise. Previous studies have determined that liver mitochondrial acetyl-CoA is increased by ketogenic insufficiency (42). In addition to ketogenesis, terminal oxidation of β-oxidation-derived acetyl-CoA in the TCA cycle is a prominent means of lipid catabolism in the liver (43). Our data show TCA cycle flux is higher in liver HMGCS2 KO mice relative to WT littermates at rest and accelerated in both genotypes during exercise. However, the greater TCA cycle flux in HMGCS2 KO mice was no longer observed after 30 minutes of treadmill running. Interestingly, we recently observed liver HMGCS2 KO mice have elevated liver TAGs despite increased TCA cycle flux when fed a ketogenic diet under sedentary conditions (14). These findings suggest that lipid disposal via the TCA cycle increases to compensate for ketogenic insufficiency. However, the TCA cycle may not be sufficient to prevent liver lipid accretion when confronted with conditions that place further demands on TCA cycle flux (i.e., an exercise challenge) and/or significantly increase lipid availability.

### Role of hepatic HMGCS2 in lipid adaptations to exercise training

Regular exercise reduces liver steatosis in both humans and experimental models (2, 19, 44–51). Given that ketogenic insufficiency induced liver lipid accumulation in response to acute exercise, we tested if loss of hepatic HMGCS2 would attenuate the effectiveness of exercise training to mitigate fatty liver. Liver-specific HMGCS2 KO mice and WT littermates remained untrained or were trained via a 6-week treadmill running protocol. Forced exercise training was performed to better facilitate the consistent implementation of key exercise stimuli including duration, intensity, and frequency (52). This was important because our studies showed that liver HMGCS2 KO mice spent more time running at higher speeds and for greater distance when provided voluntary running wheels. In congruence with previous work, liver TAGs were lower in trained WT compared to the untrained WT mice. Unexpectedly, training was equally effective at lowering liver TAGs in liver HMGCS2 KO mice as it was in WT mice. This outcome indicates that hepatic ketogenesis is not required for training to mitigate diet-induced fatty liver.

The effectiveness of training to combat fatty liver in the presence of ketogenic insufficiency is linked to multiple metabolic mechanisms. As previously discussed, TCA cycle flux is increased in liver HMGCS2 KO mice at rest. A persistent elevation in lipid disposal via the TCA cycle may aid in lowering liver TAGs in trained KO mice. Similar to liver HMGCS2 KO mice, we previously showed that the inhibition of exercise-stimulated TCA cycle flux through deletion of hepatic PCK1 in mice led to liver TAG accumulation during acute exercise, but did not compromise the lipid lowering actions of training (8). While TCA cycle flux was impaired, the liver PCK1 KO mice exhibited the delayed suppression of ketogenesis and/or ketosis upon refeeding following an acute exercise bout (8). Thus, individual mitochondrial oxidative pathways may compensate for impairments in the others over the longer timeline of exercise training.

In addition to fat disposal in hepatic mitochondrial pathways, adaptations in extrahepatic metabolism may also contribute to the prevention of fatty liver in trained mice lacking hepatic HMGCS2. We observed that training blunted the gain in body weight and fat mass in both WT and liver HMGCS2 KO mice. Repeated bouts of muscular work also enhanced markers of mitochondrial content and/or oxidation in skeletal muscle. Specifically, muscle citrate synthase activity was increased and TAGs were decreased in response to training. Our results are consistent with a recent publication that exercise training increases skeletal muscle mitochondrial respiration similarly in rats with a hepatic knockdown of HMGCS2 and WT controls (53).

Ultimately, lower adiposity and an elevation in skeletal muscle lipid oxidation in trained mice could reduce fatty acid delivery to the liver and, subsequently, limit liver lipid accretion. The decreased liver AcAc and βOHB in trained WT mice are agreeable with the hypothesis that liver lipid supply is diminished in response to training. While ketogenic insufficiency does not inhibit training from protecting against obesity, it should be noted that loss of hepatic HMGCS2 may decrease the efficiency of training to promote anti-obesity adaptations. Davis et al. reported that loss of hepatic HMGCS2 prevents a training-induced rise in energy expenditure without negatively impacting body weight or composition following 16 weeks of voluntary wheel running (54). Our results showed that training decreased the gain in body weight within two weeks in WT mice. However, the impediment in body weight gain in liver HMGCS2 KO mice was delayed until week 5 of training.

In conclusion, hepatic HMGCS2-dependent ketogenesis is important for liver lipid homeostasis during an acute non-exhaustive, exercise bout. In contrast, hepatic HMGCS2 is not required for training to treat diet-induced fatty liver. Our results suggest that the efficacy of training to lower liver fat in the absence of HMGCS2 is linked to compensatory lipid disposal via the TCA cycle and extrahepatic adaptations that limit liver lipid availability. Furthermore, these studies highlight the multifactorial and integrated mechanisms through which training reduces fatty liver.

## DATA AVAILABILITY

Data will be made available upon request.

## ACKNOWLEDGMENTS

Figure schematics were generated using BioRender.com. Time course images for indirect calorimetry experiments were created using CalR (55). The authors thank the Analytical Biochemistry shared resource at the University of Minnesota (NIH P30 CA77598) for access to GC–MS instrumentation.

## GRANTS

This research was supported by the NIH Grants DK136772 (C.C.H.), DK091538 (P.A.C.), and AG069781 (P.A.C.). The Functional Lipidomics Core at the Barshop Institute is partially supported by NIH Grants P30 AG013319 (X.H.) and P30 AG044271 (X.H.). A.F.O. was supported by the National Institutes of Health Ruth L. Kirschstein National Research Service Award T32 DK007293. E.D.Q. was supported by the National Institutes of Health Ruth L. Kirschstein National Research Service Award T32 HL166142.

## DISCLOSURES

No conflicts of interest, financial or otherwise, are declared by the authors.

## AUTHOR CONTRIBUTIONS

C.V., A.F.O., and R.E.P: Investigation, Formal analysis, Data curation, and Writing - review & editing; J.L.H., G.S.H., and H.W.: Investigation, Data curation, and Writing - review & editing; E.D.Q.: Investigation and Writing - review & editing; P.A.C. and X.H.: Resources, Writing - review & editing, and Funding acquisition; C.C.H.: Conceptualization, Investigation, Formal analysis, Data curation, Resources, Funding acquisition, Supervision, Project administration, Visualization, Writing – original draft, and Writing – review & editing.

## FIGURE CAPTIONS

**Supplemental Figure S1.**
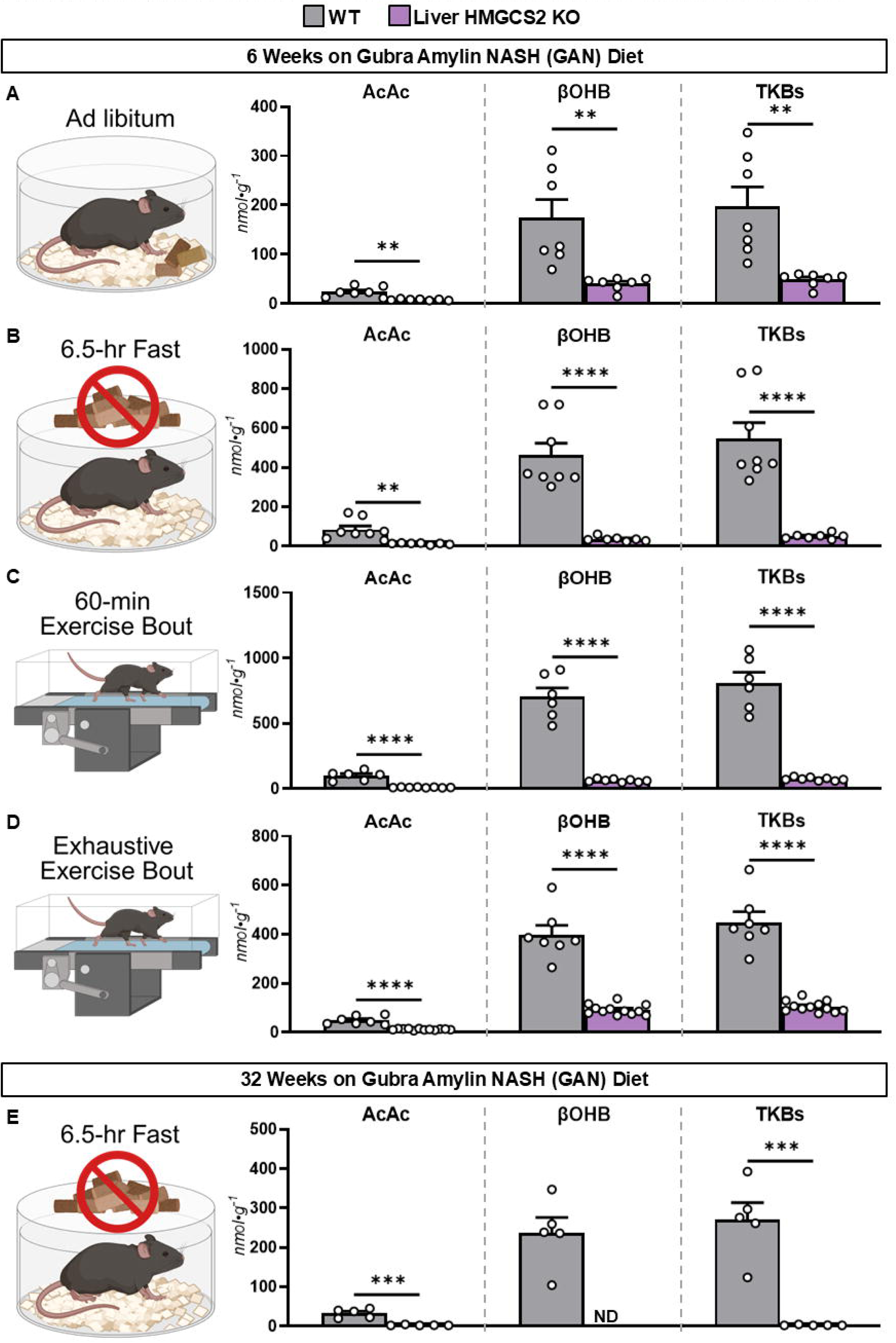
Comparison of liver ketone bodies in wild type (WT) and liver HMGCS2 knockout (KO) mice. Data points in this figure are the same as in Figure 1, however, they are displayed to promote comparison between WT and Liver HMGCS2 KO mice. Liver acetoacetate (AcAc; nmol·g^-1^), β-hydroxybutyrate (βOHB; nmol·g^-1^), and total ketone bodies (TKBs; nmol·g^-1^) in 12-week-old mice receiving a GAN diet for 6 weeks and fed ad libitum, fasted 6.5 hours, exercised for 60 minutes, or exercised to exhaustion (**A-D**; n = 6-12 per genotype). **E:** Liver AcAc (nmol·g^-1^), βOHB (nmol·g^-1^), and TKBs (nmol·g^-1^) in 38-week-old mice receiving a GAN diet for 32 weeks and fasted 6.5 hours (n = 5 per genotype). Data are mean ± SEM. **p<0.01, ***p<0.001, and ****p<0.0001. ND = not determined because concentrations below quantifiable limits.

## Notes

### Competing Interest Statement

The authors have declared no competing interest.

