## Supplemental Table 1 for "Hepatic ketogenesis supports liver lipid homeostasis during acute exercise but is not required for exercise training to mitigate liver steatosis in mice"

| Antibody | Supplier | Catalog Number | RRID | Dilution |
|---|---|---|---|---|
| β-hydroxybutyrate dehydrogenase 1 | Proteintech | 15417-1-AP | RRID:AB_2274683 | 1:1000 |
| Glucose-6-phosphatase | Proteintech | 22169-1-AP | RRID:AB_2879015 | 1:1000 |
| Glycogen phosphorylase | Proteintech | 15851-1-AP | RRID:AB_2175014 | 1:1000 |
| Phospho-glycogen phosphatase (Ser15) | Abcam | ab227043 | N/A | 1:1000 |
| Phospho-glycogen synthase (Ser641) | Cell Signaling Technologies | 3891 | RRID:AB_2116390 | 1:1000 |
| Glycogen synthase 2 | Proteintech | 22371-1-AP | RRID:AB_2879091 | 1:1000 |
| Phosphoenolpyruvate carboxykinase 1 | Proteintech | 16754-1-AP | RRID:AB_2160031 | 1:1000 |
| 3-hydroxymethylglutaryl-CoA synthase 2 | Cell signaling Technologies | 20940 | RRID:AB_2798853 | 1:3000 |
| Anti-rabbit IgG | Cell Signaling Technologies | 7074 | RRID:AB_2099233 | 1:5000 |

**Supplemental Table S1**. **List of antibodies.**
